# Different trajectories of polyploidization shape the genomic landscape of the *Brettanomyces bruxellensis* yeast species

**DOI:** 10.1101/2021.02.11.430840

**Authors:** Chris Eberlein, Omar Abou Saada, Anne Friedrich, Warren Albertin, Joseph Schacherer

## Abstract

Polyploidization events are observed across the tree of life and occurred in many fungi, plant and animal species. Polyploidy is thought to be an important source of speciation and tumorigenesis. However, the origins of polyploid populations are not always clear and little is known about the precise nature and structure of their complex genome. Using a long-read sequencing strategy, we sequenced a large number of isolates from the *Brettanomyces bruxellensis* yeast species, which is found in anthropized environments (*e.g.* beer, contaminant of wine, kombucha and ethanol production) and characterized by several polyploid subpopulations. To reconstruct the polyploid genomes, we phased them by using different strategies and we found that each subpopulation had a unique polyploidization history with distinct trajectories. The polyploid genomes contain either genetically closely related (with a genetic divergence < 1%) or diverged copies (> 3%), indicating auto- as well as allopolyploidization events. These latest events have occurred independently with a specific and unique donor in each of the polyploid subpopulations, and exclude the known *Brettanomyces* sister species as possible donors. Finally, loss of heterozygosity events have shaped the structure of these polyploid genomes and underline their dynamic. Overall, our study highlights the multiplicity of the trajectories leading to polyploid genomes within a same species.

## Introduction

Polyploidy, a state in which organisms carry three or more full sets of chromosomes, is a phenomenon that can be observed throughout plant, animal and fungi species. Polyploidization has gained in interests because of its tremendous effects in the evolution of species or its involvement in cancerogenesis (Adams and Wendel 2005; Gregory and Mable 2005; Gjelsvik et al. 2018). The most obvious and probably well studied polyploidy events in the tree of life are the Whole Genome Duplication (WGD) events, which are usually followed by subsequent and massive diversification. One example is the series of two ancient WGD events that occurred in the lineage leading to the ancestor of all vertebrates ~450 million years ago, and have significantly contributed to the successive evolution of 60,000 extant species (Dephal and Boore 2005; Sacerdot et al. 2018).

While the effects of polyploidization on the organisms are versatile, there are also different types of polyploids (Comai 2005; Fox et al. 2020). The doubling of one’s own genome or the merging of genomes coming from individuals of the same species would lead to multiple genomic copies of identical or similar descent, and defines the mechanisms of autopolyploidization. However, interspecific hybridization associated with genome doubling would cause the acquisition of additional full chromosomal sets harbouring higher genetic variation and defines allopolyploidization. While it is well observed that a polyploid state causes genomic conflicts, leads to genome instability, or reduces gamete formation, on the contrary, genomic reorganization can promote diversification from the additional genomic information (Mayer and Aguilera 1990; Wood et al. 2009; Van de Peer et al. 2017). Therefore, polyploidy play a predominant role in bursts of adaptive divergence and speciation (Leitch and Leitch 2008; Soltis et al. 2015). Polyploidy can be beneficial under certain environmental circumstances and increases the potential for adaptability, taking advantages of evolutionary innovations from neo- and subfunctionalization of duplicated genes (Sanchez-Perez et al. 2008; Eberlein et al. 2018). Environmental changes that require a fast adaptation for example can trigger the prevalence of polyploids, which at least for short-terms, can have an adaptive advantage from genomic flexibility rather than simply being the “dead-end”.

Some taxa are believed to be more stable to polyploid states than others. These are known to be found frequently among plants, which in contrast to animals are characterized by a development that seems to be more robust to genomic perturbations (Orr 1990). Studies suggest that up to 70% of flowering plants originate from polyploid ancestors, putting it as a major contributor in the evolution of species (Masterson 1994; Levin 2004). But it is also suspected in animals that polyploidization plays an even more prevalent role than currently shown, but limited to the analytic tools and effort detecting them. While animals are characterized by less stable polyploids, it is well acknowledged that most of the vertebrate species originate from ancient polyploidization events too (Gregory and Mable 2005; Wertheim et al. 2013). At the same time, polyploidy is also increasingly observed in single-cell organisms such as yeasts (Bellon et al. 2011; Krogerus et al. 2015; Peter at al. 2018), suggesting that this state can be used as a rapid response to ecologically or human-made changes in anthropogenically-used environments, coevolution or enable invasions by the acquisition or holding of additional full sets of chromosomes (Te Beest et al. 2012; Bertier et al. 2013; Morrow and Fraser 2013; Van de Peer et al. 2017). In the lineage leading to *Saccharomyces cerevisiae,* a hybridization event between two ancestral species have been followed by subsequent WGD, a mean by which, in the subsequent process of extensive genome reorganization, high fertility could be retained (Seoighe and Wolfe 1998; Gerstein and Otto 2009; Gordon et al. 2009; Marcet-Houben and Gabaldon 2015). Moreover, the prevalence of polyploidy, currently observed in *S. cerevisiae* is approximately 11.5%, as shown in a recent study of 1,011 whole-genome sequenced isolates (Peter et al. 2018). Polyploids are particularly enriched in subpopulations associated with the production of beer or bread, highlighting that yeast domestication most likely triggered the selection of polyploids displaying desired requirements in industrial settings.

With polyploidization being recognized as a ubiquitous mechanism in nature with almost unpredictable consequences in terms of genomic conflicts or adaptability, we are still at the beginning to fully resolve and understand the genomic architecture of natural polyploid populations, their prevalence, and trajectories especially within the same species. The genomic era has accelerated the research on polyploid and hybrid genomes through the access of long-read sequencing data. However, biggest challenges are still the correct phasing of haplotypes, to separate the different set of chromosomes without any prior knowledge about ploidy and levels of genetic variation between genomic copies. Here, we focused on the *Brettanomyces bruxellensis* yeast species, a genetically diverse species with different subpopulations of various levels of ploidy allowing to shed light into several questions related to polyploidization. As seen for other yeasts of the *Saccharomycotina* subphylum, the link between ecological origin and genetic differentiation for the different *B. bruxellensis* clades is primarily supposed to be driven by its anthropogenic influences (De Barros Pita et al. 2011; Avramova et al. 2019). Multiple genetically distinct subpopulations (clusters) correspond to different ecological niches, respectively wine, beer, tequila/bioethanol, kombucha and soft drinks (Avramova et al. 2018).

To study their genomic complexity and to allow a detailed view on their genomic architecture for the first time, we sequenced a subset of 71 *B. bruxellensis* strains from different subpopulations with long and short reads sequencing strategies. By using two complex phasing strategies, our results highlighted that polyploidy, even within the same species, can follow different trajectories, respectively auto- and allopolyploidization. We suggest the need to applying phasing algorithm to species with polyploid genomes as a basis for further resolving the link between different trajectories of polyploidy and their phenotypic consequences in an ecological diverse setting.

## RESULTS

### Conserved clusters of polyploid isolates

*Brettanomyces bruxellensis* is known as a diverse species with genetically and ecologically distinct clusters, and various levels of ploidy (Avramova et al. 2018, Gounot et al. 2020, Colomer et al. 2020). To dissect the genomic architecture and to further understand the origin as well as the trajectories of recently described polyploid groups, we selected 71 strains with 51 coming from subpopulations defined as polyploids (Avramova et al. 2018) (Figure 1A; Supplementary Table S1). Most of the strains were isolated in Europe and originate from different ecological origins: beer (n=25), wine (n=36), tequila/bioethanol (n=7) and kombucha (n=3).

**Figure 1.**
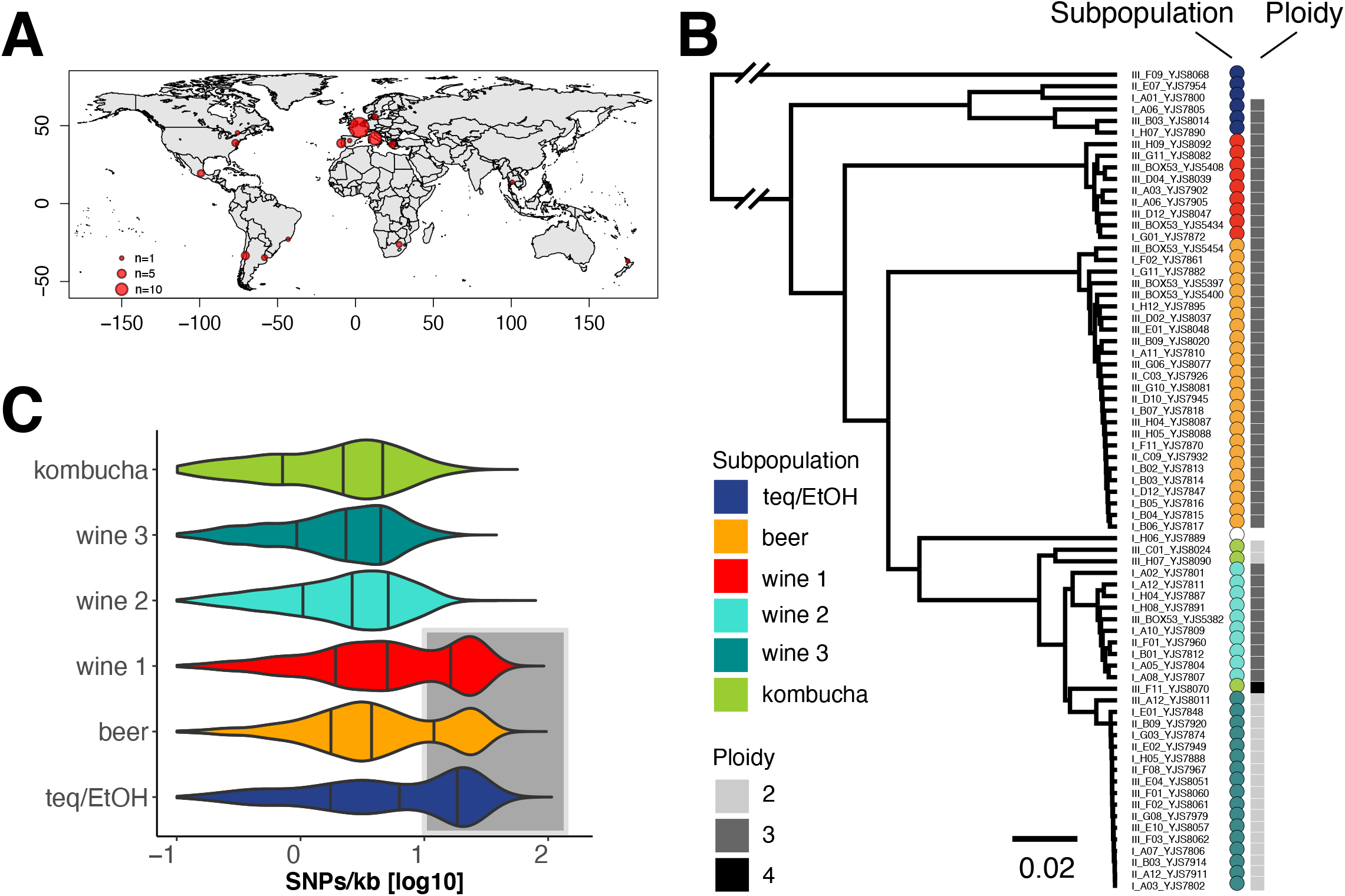
Ploidy and intra-genomic variation. **A** Strain collection. The 71 sequenced strains come from the collection of 1,500 isolates (Avramova et al. 2018) and were isolated in different regions worldwide, where they are associated with anthropized environments such as tequila/bioethanol, beer, wine and kombucha production. **B** Genetic relationship and ploidy level. The sequenced strains, here clustered based on Illumina short read sequencing data (75PE), separate in six genetically distinct subpopulations, namely tequila/bioethanol (teq/EtOH), beer, wine (1-3) and kombucha (based on 24,313 genome-wide distributed variants). Forty-eight strains were detected as triploids (69%) coming from five of the six subpopulations: teq/EtOH, beer, wine (1,2) and kombucha (inferred from genome-wide allele frequencies). **C** Genetic diversity within clades inferred from long-read sequencing data. The three subpopulations teq/EtOH (n=5), beer (n=22) and wine 1 (n=7) harbour strains with two clusters of reads bearing low and high genetic variation (underlaid in grey) compared to the reference genome *Brettanomyces bruxellensis* (Fournier et al. 2017). The subpopulation wine 2 (n=9), although being polyploid (**B**), lacks genomic regions with high genetic variation to the reference genome. The three lines within each distribution show the 25%, 50% and 75% quartiles.

To have a deep insight into the population structure and ploidy levels, we first sequenced the 71 genomes using a whole genome Illumina short-read sequencing strategy with a 16.9-fold mean coverage. Using this dataset, we sampled 24,313 genetic variants evenly distributed across the genome and performed a phylogenetic analysis (Figure 1B). All the 71 strains were clustered into six well-defined lineages, which correlate with environmental niche, as previously reported (Avramova et al. 2018, Gounot et al. 2020, Colomer et al. 2020). We then assigned ploidy state to each of the strains using SNP frequency distributions within the sequence reads. We categorized their level of ploidy as either being diploid (with SNPs at allele frequencies at 0.5 and 1), triploid (with SNPs at allele frequencies at 0.33 and 0.67) or tetraploid (with SNPs at allele frequencies at 0.25, 0.5, 0.75 and 1) (Supplementary Figure S1A). We found that the level of ploidy is conserved within, but varies across subpopulations (Figure 1B). The wine 1, wine 2 and beer subpopulations are triploid while the wine 3 and kombucha subpopulations are diploid. The exception is a single kombucha strains being tetraploid (Supplementary Figure S1A). The teq/EtOH clade harbours three triploid strains while the ploidy could not have been assigned to the three other isolates (Supplementary Figure S1A). For another single strain (I_H06_YJS7889), we could neither identify its ploidy, nor assign it to one of the six subpopulations. To exclude that aneuploidies are causing the non-assignment of ploidy levels for the four strains, we looked at read coverage across their genome to identify regions that are absent or present in multiple copies (Supplementary Figure S1B-C). We showed that the coverage is steady and that these strains do not contain regions with varying coverage explaining our results.

Overall, we can highlight that the level of ploidy is conserved within genetically diverged subpopulations, but not across them. We showed that the teq/EtOH strains are the most diverse subpopulation and confirmed previous data that additionally suggested this subpopulation as the oldest of the different *B. bruxellensis* clades (Colomer et al. 2020). The teq/EtOH strains stand in contrast to other subpopulations like the wine 3, which show the lowest degree of genetic variation and therefore might hint towards a single ancestral origin with a recent expansion.

### Strategies used to phase the *B. bruxellensis* polyploid genomes

In order to resolve the genomic structure of polyploid isolates, we sequenced the genomes of the 71 strains using the Oxford Nanopore sequencing strategy. Long-read sequencing has become the strategy of choice to best resolve structural variation and build high quality *de novo* reference assemblies. The difficulty of resolving polyploid genomes, however, lies especially in the attempt to distinguish between the different haplotypes, which are present as independent genomic copies within the same genomes. While *de novo* assemblers are not capable of fully separating different haplotypes, providing instead collapsed haplotypes, several alignment-based algorithms have been developed recently to cope with the genomic architecture of polyploid genomes (Schrinner et al. 2020, Abou Saada et al. 2020, Shaw and Yu 2020). They all aim to phase haplotypes into independent entities, but they vary in performance depending on factors such as ploidy, coverage, and the level of genetic divergence between genomic copies of the polyploid genomes.

To properly phase our polyploid genomes, we sought to apply different strategies depending on the level of divergence of the copies constituting these genomes. Reasons to believe that there are different levels of variation have been given by previous studies, which indicated that at least two single polyploid strains from the wine 1 and beer subpopulations have likely experienced polyploidization events by having an additional genomic copy of high genetic variation (Borneman et al. 2014).

To estimate the genetic divergence, we hence aligned the long reads of each strain to the *B. bruxellensis* reference genome (Fournier et al 2017; Supplementary Figure S2A). We could identify three subpopulations (teq/EtOH, beer and wine 1) for which the genetic variation resolves in a bimodal distribution with a low as well as a high genetic variation level cluster of reads (Figure 1C). These subpopulations stand in contrast to the other three subpopulations (wine 2, wine 3 and kombucha), which solely comprise low genetic divergent reads. Strikingly, wine 2, is the only polyploid subpopulation which bears reads with only low genetic diversity.

With the two types of polyploid subpopulations exhibiting either low or high genomic variation, we further applied two different phasing strategies to study their genomic architecture.

(1) To resolve the origin of the genetic diversity, and to determine if the cluster with reads of high genetic variation complements an additional genomic copy, we separated the long sequencing reads into distinct clusters based on their diversity level. We clustered reads with peaks of low genetic variation at 2 SNPs per kb and high genetic variation exhibiting 24.4 SNPs per kb (Supplementary Figure S2A-B). Reads between the two distribution (*i.e.* with a variation between 10 and 14 SNPs per kb) were ignored to avoid the assignment of reads to the wrong cluster (Supplementary Figure S2B). Based on these sets of reads, we generated *de novo* assemblies in order to have the phased copies of the polyploid genomes.

(2) The low genetic variation observed in the polyploid wine 2 subpopulation did not allow us to separate reads based on their genetic divergence (Figure 1C). Consequently, we used nPhase, a phasing algorithm that we recently developed (Abou Saada et al. 2020). Briefly, nPhase resolves the genome into distinct haplotypes and provides accurate and contiguous haplotype predictions using short and long read sequencing data without any prior information of the true ploidy. It accurately identifies heterozygous positions using highly accurate short reads and clusters long reads into haplotypes based on the presence of similar heterozygous SNP profiles (Abou Saada et al. 2020).

### Genomic architecture of the polyploid wine 2 subpopulation

We applied the nPhase phasing algorithm to the sequenced genomes of the wine 2 subpopulation, exhibiting a low intra-genomic variation. We focused on six of the ten strains for which we had high quality long and short read sequencing data, allowing us to phase their genome properly into independent haplotigs (Figure 2; Supplementary Figure S3A).

**Figure 2.**
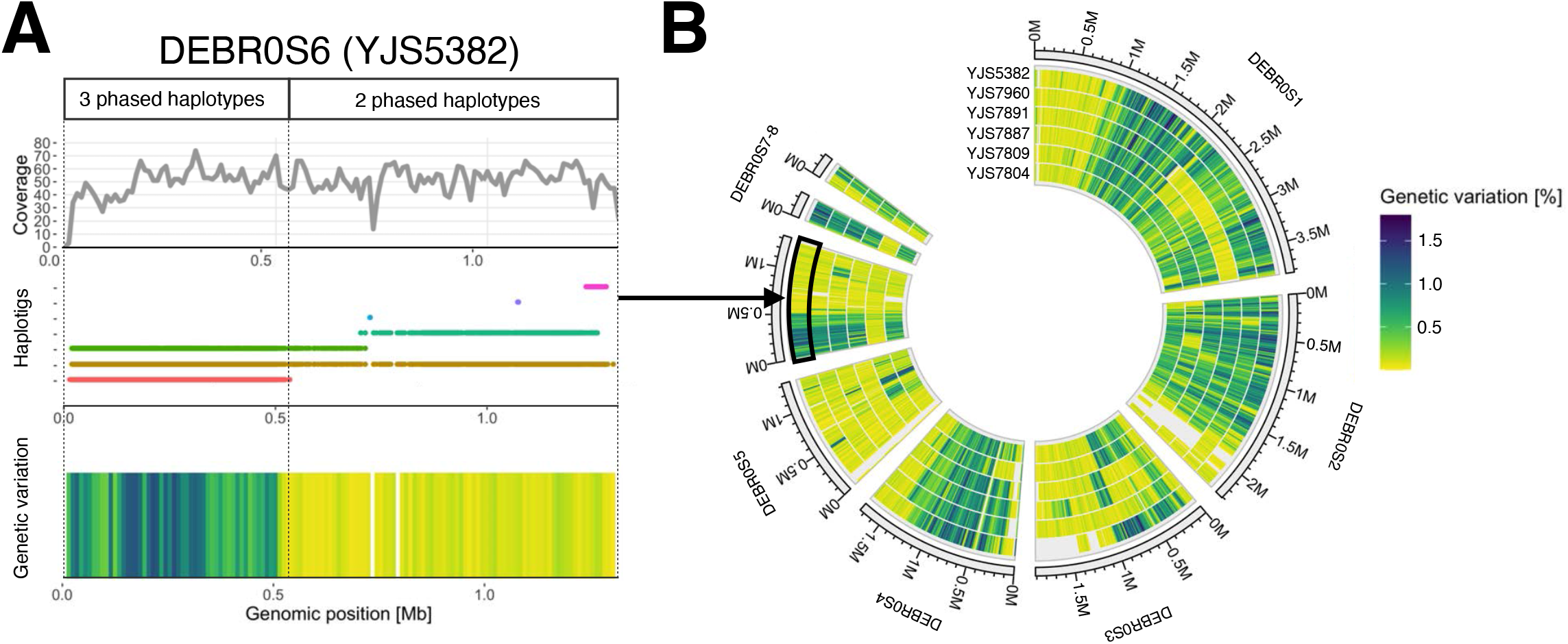
Autopolyploidization for the wine 2 subpopulation. **A** Separation of haplotypes. Phasing the genomes of strains from the polyploid wine 2 subpopulation resolves the generally low intra-genomic variation into haplotigs along the genome. The presence of two haplotypes resolves in lower genetic variation as it does when three haplotigs are present at a given position. Maximal genetic variation between haplotypes increases from 0.93 % to 1.79 % with the presence of a third phased haplotype. To control that the variation in genetic difference are not artefacts coming from a variable coverage along these regions, the genome-wide coverage was calculated. The coverage is consistent across regions that harbour either two or three phased haplotypes. **B** Conserved patterns of phased haplotypes along the genomes of six strains of the wine 2 subpopulation. Having either two or three phased haplotypes at a site is conserved among different strains from the same subpopulation.

We observed that the chromosomes are phased into regions underlying in most cases two or three haplotigs (Supplementary Figure S3A). Some regions bear multiple and often small haplotigs, and underline the complexity in phasing polyploid genomes with haplotypes that reflect low genetic similarity. In addition, the level of genetic divergence varies along the genomes of the six strains. Whenever nPhase resolves a region into two haplotigs, the genetic variation in these regions is lower compared to regions where it distinguishes between three haplotigs (Figure 2A). Here, the highest genetic variation in the presence of two haplotigs is 0.93%, while on average it is as low as 0.09% (Figure 2B, Supplementary Figure S3B). In the presence of a third phased haplotig, the genetic variation can be as high as 1.79% with an average genetic variation of 0.54% (Figure 2B, Supplementary Figure S3C). Consistent coverage levels support the hypothesis that the prediction of only two haplotypes is not due to the absence of a third copy for part of the chromosome. (Figure 2A). Therefore, while the differentiation of three haplotigs underlines the existence of three genetically different genomic copies at that site, the phasing resolving into two haplotigs represent a region with two identical haplotypes plus the existence of genetically different copy.

Further, we can show the presence of conserved regions in all six strains that are characterized by the presence/absence of a third phased haplotype (Figure 2B; Supplementary Figure S3B-C). Some regions, for example the first 1 Mb on chromosome 1, are characterised by two identical copies and a non-identical copy, resolving into two phased haplotypes. This region is followed by another 1 Mb region, which resolved into three haplotypes in all six strains. An explanation for the alternation of such regions phased into two or three haplotypes is the occurrence of loss of heterozygosity (LOH) events. LOH are characterized by the removal of polymorphic markers that distinguish different genomic copies in diploid or polyploid individuals and consequently reduced the genetic variation. nPhase outputs only unique haplotypes present in the data, irrespective if one haplotype contains twice the number of reads as the other, and therefore indicates the occurrence of LOH events in the genome. Moreover, the existence of the conserved regions of LOH events among the six strains can hint at hotspots for LOH events. Such hotspots have been shown in other species such as *S. cerevisiae* (Peter et al. 2018), where they frequently cause the removal of genetic variation. Alternatively, this conserved pattern could also hint towards a recent common ancestor. But with these strains being isolated in different countries of two continents (Supplementary Table S1), this explanation is less likely.

Overall, the utilization of long and short read sequences in combination with complex phasing strategies enables to decipher the genomic structure of polyploid genomes of low genetic variation and allows to study its dynamic. Further, for the wine 2 subpopulation, the only polyploid clade with a low intra-genomic variation, the genomes of six strains revealed conserved regions having undergone LOH events.

### Three polyploid clades contain a genetically diverged genomic copy

Next, we focused on the triploid genomes of the teq/EtOH, beer and wine 1 subpopulations, which exhibit genetically very heterogeneous genomes. To enable comparative analyses, we first separated reads based on their genetic divergence compared to the reference genome (Supplementary Figure S2A). We clustered reads from the bimodal distribution with reads bearing low genetic variation (peak at 2 SNPs per kb) and reads with high genetic variation (peak at 24.4 SNPs per kb) (Supplementary Figure S2B). As previously mentioned, reads with a variation between 10 and 14 SNPs per kb were ignored to avoid assigning reads to the wrong cluster. The determination of the ratio between the number of reads with a low genetic variation and the total coverage (reads with low and high genetic variation) within 10 kb windows across the genome allowed us to determine the average genomic ploidy level of each strain at a given genomic position (Supplementary Figure S4A). We identified that the three groups (teq/EtOH, beer and wine 1) contained two genomic copies with low genetic variation and a single genomic copy that exhibit a high genetic divergence (or vice versa), which on average complemented to 3n genome-wide (Supplementary Figure S4B).

The fact that the beer and wine 1 subpopulations contain isolates with higher genetic variation compared to the reference genome was already shown previously for a single strain from each subpopulation (Borneman et al. 2014). We can, for the first time, highlight that this phenomenon of having a genetically different genomic copy within these subpopulations is frequent and conserved. Additionally, while previous analyses have underpinned the prevalence of polyploid strains in the teq/EtOH subpopulation, we can also show that teq/EtOH strains contain a genomic copy as genetically different to the reference genome of *B. bruxellensis* as in beer and wine 1 isolates.

To allow comparative studies between the different genomic copies among these genomes as well as with genomes from the other subpopulations, we first performed *de novo* genome assemblies using *SMARTdenovo* (Liu et al. 2020). We generated independent *de novo* assemblies for the different genomic copies, respectively reads bearing low or high genetic variation for all strains from the three subpopulations (teq/EtOH, beer and wine 1) (Supplementary Figure S5; Supplementary Table S2). Then, to base the comparative analysis on reliable Illumina short-reads sequencing data, we used group-specific reference genomes by concatenating *de novo* assemblies from reads containing low and high genomic variation (See: Material and Methods). By performing a competitive mapping, we separated the short reads for each strain from the three groups (teq/EtOH, beer and wine 1) (Supplementary Figure S5). Then, we aligned the reads back to the *B. bruxellensis* reference genome. For strains that were either diploid, or polyploid without reads with high genetic variation compared to the reference genome, we aligned short read sequences directly to the reference genome (wine 2, wine 3 and kombucha subpopulations).

We determined the genetic diversity of the 71 strains by performing a principal component analysis (Figure 3A). By looking at the first two principal components explaining 53.7% of the variation from 24,110 sampled genome-wide distributed SNPs, we can show that the genomic copies with high genetic variation (‘High’) of 40 strains from the teq/EtOH, beer and wine 1 subpopulations are clearly distinct from the genomic copies with low genetic variation (‘Low’), and cluster group-specific.

**Figure 3.**
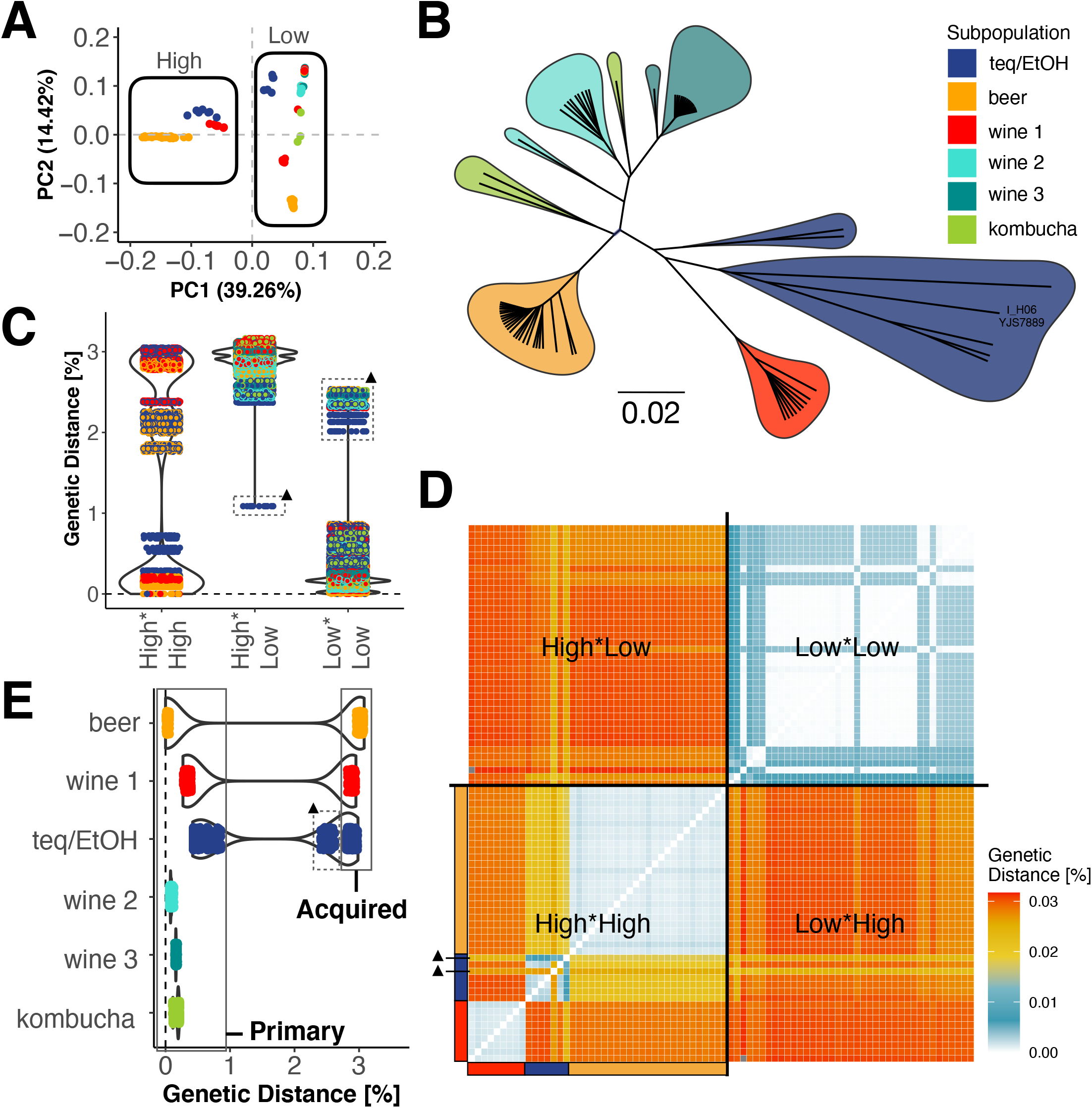
Three independent interspecific hybridization events. **A** Three distinct clusters of genomic copies with high genetic variation. A principal component analysis shows that the genomic copies with high genetic variation to the reference genome in the subpopulations teq/EtOH, beer and wine 1 are not only different to the genomic copy with low genetic variation, but are also genetically distinct between subpopulations (based on 24,110 genome-wide distributed SNPs). **B** Phylogenetic relationship from reads with low genetic variation to the reference genome. The genomic copies with low genetic variation are different between the six subpopulations teq/EtOH, beer, wine 1-3 and kombucha, which group according to their ecological origin (based on 24,110 genome-wide distributed SNPs). **C-E** Pairwise genetic diversity between genomic copies from imputed whole-genome sequences. **C** The pairwise genetic analysis shows that the genomic copy with high levels of intra-genomic variation within subpopulations is less than 1% (average of 0.13%). Between subpopulations, it varies between 1.76% and 3.05% (High*High). The genetic distance between the genomic copies with low and high levels of variation is on average 2.92% (High*Low). Genetic distances between genomic copies of low intra-genomic variation are generally below 0.9% (Low*Low). The strain III_F09_YJS8068 is an outlier (black triangle) as it has the closest distance between its own two different genomic copies with 1.1% (High*Low). It also has the highest variation of its low intra-genomic copy to the other low-intra genomic copies of other strains (Low*Low; >2%). **D** Heatmap showing the genetic distance between genomic copies of 40 polyploid individuals. The only strains whose genomic copy with low intra-genomic variation is most close to its genomic copy with high intra-genomic variation is III_F09_YJS8068 (black triangle), which clusters with all other genomic copies of high intra-genomic variation. **E** Acquired genomic copy of unknown origin. The genomic copies with low genetic variation were assigned as the primary genomic copies present in all individuals (2n-4n), while the genomic copies with high genetic variation were assigned as acquired genomic copies, only present in 40 polyploids of the subpopulations teq/EtOH, beer and wine 1. Pairwise genetic analysis with the primary genome of the beer subpopulation as a reference show a clear gap between the genetic variation that defines the primary and the acquired genome. The primary genome of the beer subpopulation is similarly distant to its own acquired genomic copy as well as to the acquired genomic copies of the other two polyploid groups wine 1 and teq/EtOH. The two genetics cluster beyond the 0.9% for the teq/EtOH comprise not only the pairwise comparison with the acquired genomic copies. For the strain III_F09_YJS8068 (black triangle), the primary genomic copy is nearly as genetically distinct as the acquired genomic copy. Genetic distances were calculated pairwise per chromosome and then average per genome (JC69).

We then checked for the genetic relationship of the genomic copies with only low genomic variation, since such genomic copies were present in all of the 71 strains (Figure 3B). We can show that the strains cluster in the six subpopulations as previously seen using the raw Illumina data (Figure 1B). The single strain I_H06_YJS7889, initially unable to be associated with a subpopulation, now clusters with other teq/EtOH strains.

### The acquired divergent copies highlight clade specific allopolyploidy events

To study the origin of the genetically divergent copies present in the three subpopulations, we imputed whole-genome fasta-alignment files for every individual. We calculated pairwise genetic distances and found that the divergence between these copies within the subpopulations was 0.13% on average (Figure 3C; High*High). By contrast, the genetic divergence of these copies across the subpopulations was 2.59% on average, ranging from 1.76% to 3.05%.

When comparing the genetic distance between the low and highly diverged genomic copies across all the genomes, we observed that the genetic distance is 2.92% on average (Figure 3C; High*Low). With 3.16%, the largest genetic distance can be seen between the wine 1 and kombucha subpopulations. The single outlier strain is III_F09_YJS8068 (teq/EtOH) and has the closest genetic distance between its own two different genomic copies, with about 1.1%. With more than 2%, III_F09_YJS8068’s low variation genome is also the most distant one to all other low variation genomes (Figure 3C). The other genomes bearing low genetic variation are less than 1% diverged. In fact, using the representation of pairwise distances in the heatmap format reassert the three genetically distinct entities of the genomic copies with high genetic variation (Figure 3D), while the genomic copies with low genetic variation being more similar (Figure 3D). Pairwise comparison using the low variable genomic copies of the beer clade as a reference confirm that inter-clade transfer of genomic copies can be excluded as a potential cause in the acquisition of additional genomic copies with high genetic variation in the three polyploid subpopulations (Figure 3E), and underpin the presence of additional and unrelated copies in these three groups.

Since there is a conservation of a closely related diploid genome across the isolates of the species, we define this part as the primary genome of *B. bruxellensis* (Figure 3E). It is present in all of the strains and harbours a genetic variation of less than 1% to the reference genome. The exception is the strain III_F09_YJS8068 strain, which groups within the teq/EtOH subpopulation, and, which has a primary genome with a minimum genetic distance of 2.01% and maximum genetic distance of 2.53% to the other primary genomes. In addition to these primary genomic copies, a highly divergent copy is present in three groups (teq/EtOH, beer and wine 1 subpopulations) and was defined as a new or ‘acquired’ genomic copy (Figure 3E). While it clearly exceeds the genetic variation of the primary genome, the acquired genomic copies open the discussion where they originate from and if they have been acquired due to interspecies hybridization.

To test whether the additional copies have been acquired from sister species as part of the genus *Brettanomyces*, we sequenced and generated *de novo* genome assemblies for four of the sister species: *B. anomala, B. nanus, B. custerianus* and *B. acidodurans* (Supplementary Table S3; Supplementary Figure S6A-D). While we were able to show collinearity between the acquired copies and the reference genome *B. bruxellensis* (Supplementary Figure S6E-F), the genomes of the sister species to *B. bruxellensis* were too dissimilar to retain any correlation using the same parameters. Only lowering the parameters were able to show a correlation and suggest less synteny paired with high genetic differentiation (Supplementary Figure S6G), as already shown by Roach and Bornemann (2020). With a genetic divergence of 2.5-3% between the acquired to the primary genomic copies, however, it seems unlikely that sister species with a genetic similarity of less than 77% have been involved in the acquisition of the additional genomic copies (Roach and Bornemann 2020).

Overall, we have shown that the triploid genomes of the wine 1, teq/EtOH, and beer subpopulation are composed of a primary part, which is common to every *B. bruxellensis* isolates, as well as a newly acquired divergent copy. These results strongly suggest that these events must have occurred independently with closer, so far unknown and far related isolates that we would, according to the genetic distance of ~3%, define as different species to *B. bruxellensis.*

### LOH events shaping the genomic landscape of interspecific hybrids

Hybrid genomes are dynamic entities with LOH events playing an important role in their evolution (Smukowski et al. 2017; Lancester et al. 2020). As already seen for the triploid genomes of the wine 2 subpopulation, these events can cause the removal of genetic variation along the genomes in a conserved manner (Figure 2B). Moreover, these events would resolve in a difference of genomic content from the parental genomes. When preparing *de novo* assemblies from reads with either high or low intra-genomic variation, we observed significantly shorter assemblies (median 9.1 Mb) for the genomic copies harbouring high intra-genomic variation (p-value = 1.3e-10; Supplementary Figure S7A). In fact, strains from different subpopulations showed a trend in which even assembly size seemed to be not only different but also conserved between subpopulations (Supplementary Figure S7B). Therefore, with the significantly shorter *de novo* assemblies for the acquired genomic copies, we hypothesized that these polyploid genomes with heterogenous levels of genetic variation have undergone LOH events as well.

To check for LOH events along the polyploid genomes, we looked at the coverage from reads belonging to the primary and acquired genome, determined if they are complementary to the total coverage, and analysed their proportion to the total coverage. Here, we used the coverage from the short reads, aligned to the reference genome *B. bruxellensis* and prior separated using competitive mapping (Supplementary Figure S5) along the chromosomes to check for reciprocal shifts in coverage (Supplementary Figure S8A-C). We can show that regions lacking reads aligned to the acquired genomic copy reveal an increase in coverage at the primary genome complementing the total coverage. On the other hand, this also appears to be the case of several regions of the primary genome, where aligned reads represent only a single genomic copy (1/3 of the total coverage), while the acquired genomic copy appears to be present in two copies (2/3 of the total coverage). These results confirm the reorganization of polyploid genome through LOH events in the subpopulations teq/EtOH, beer and wine 1.

Then, we used the primary genome as a reference and determine how many copies are present throughout the genome within the polyploid strains. We calculated its ratio per 10 kb non-sliding windows to the total coverage to assess its proportion. Our results show that the polyploid have undergone massive LOH events (Figure 4). Most regions appear to have been lost/gained within a subpopulation-specific pattern. On the chromosome 1 for example, the beer and wine 1 subpopulations lack a significant part of the acquired genomic copy (1.7 Mb for beer strains and 1.2 Mb for wine 1 strains). Other often small events, are private to single strains.

**Figure 4.**
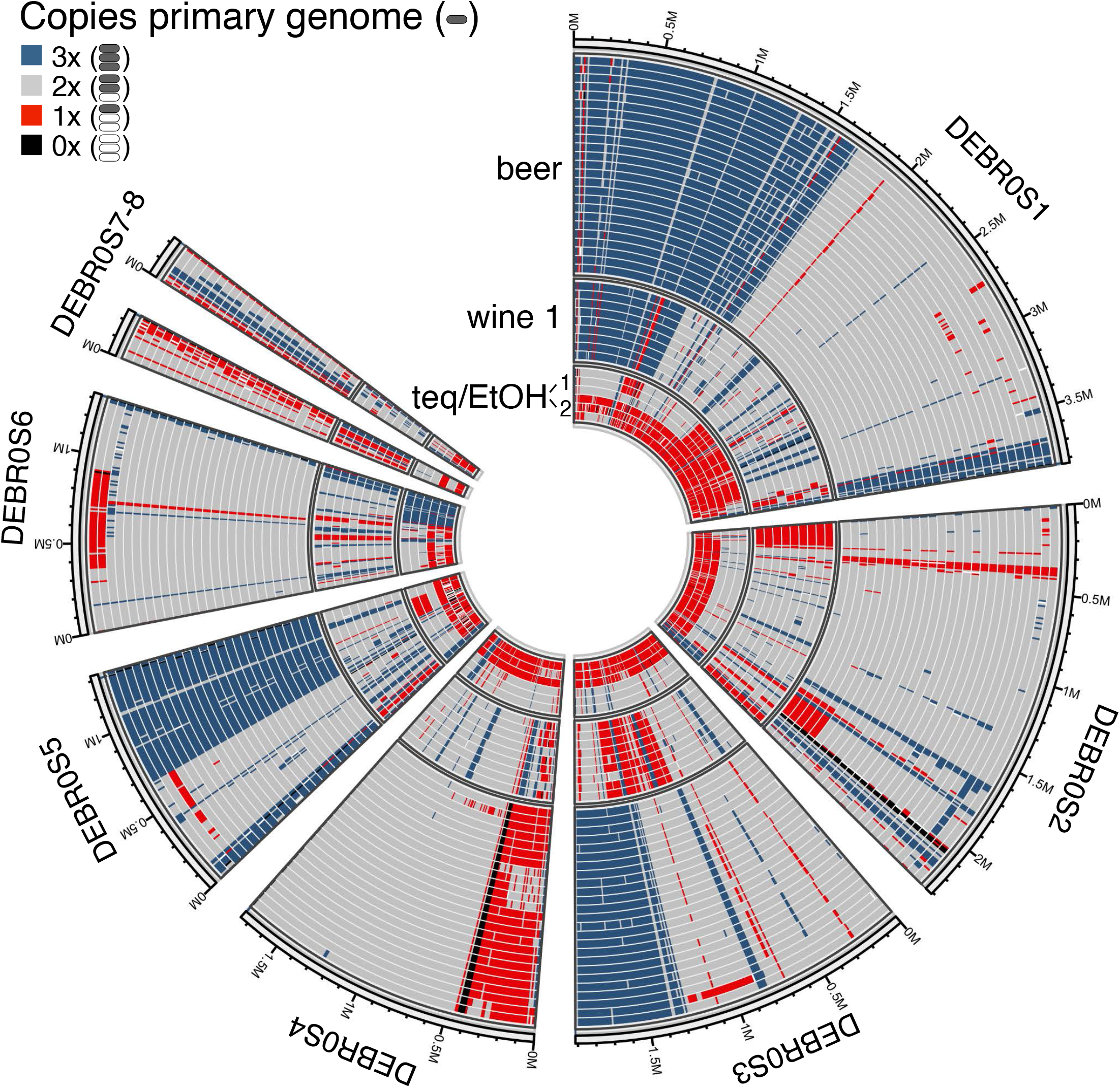
Dynamic genomic landscape of polyploid strains. The polyploid genomes of the three subpopulations beer, wine 1 and teq/EtOH underlie massive modifications through LOH events. The primary genome (low genetic variation) was used as a reference. Conserved patterns of modified regions for the primary genome were identified by determining the gain or loss of its copies in each strain, here varying between three (3x) and zero (0x). Only a few modified regions are unique to single or few strains. There are no common regions, which show the same patterns across subpopulations. The teq/EtOH subpopulation shows a division into two clusters, each consistent of three individuals. The ploidy level was estimated in 10 kb windows.

Next, we determined if the parts absent of the acquired genomic copy in the three subpopulations can complement the smaller *de novo* genome assemblies (Supplementary Figure S7). For the beer strains, we calculated that LOH events have caused the loss of 26.6% of regions in average from the acquired copy (Supplementary Figure S8A). This makes 9.54 Mb of the acquired genomic copy still kept, which is close in size to the *de novo* assembly of 8.9 Mb (Supplementary Figure S7B). For the wine 1 subpopulation, 22.3% of the acquired genomic copy is lost on average, making 10.1 Mb still being present (Supplementary Figure S8B). Here, the *de novo* assembly size (10.1 Mb) matches exactly the number of retained regions in our analysis (Supplementary Figure S7B).

The teq/EtOH strains however, show a pattern of loss/gain of genomic regions from the primary genome that enables the distinction of two subgroups, denoted as teq/EtOH 1 and teq/EtOH 2. The teq/EtOH 2 has almost entirely lost the second copy of the primary genome, being replaced by a second copy of the acquired genome (Figure 4). Both subclades have lost 10.5% of the acquired genomic copy on average (12.1% for teq/EtOH 1 and 8.8% for teq/EtOH 2). The average of both (11.64 Mb) is comparable to the average *de novo* assembly size of 10.7 Mb.

The conserved patterns of LOH within each subpopulation opens the discussion if these patterns are the consequence of adaptation, random processes, or point at a recent shared ancestry. In the evolution of species, polyploidy has been shown to potentially play an important role in the acquisition of new traits or the amplification of already existing traits in the context of the acquisition of resistances (Jackson and Tinsley 2003; Augustune et al. 2013), interactions (Thompson et al. 2004; Těšitelová et al. 2013), coping with changing environments (Selmecki et al. 2015), or the occupation of novel ecological niches (Wani et al. 2018). Further, the different environments where *B. bruxellensis’* polyploid subpopulations are associated with, respectively bioethanol production, wine or beer fermentation, are harsh environments and require different characteristics from the strains as a high tolerance against alcohol and acidity, for example.

To justify the conserved pattern per group from an adaptive evolutionary perspective, we checked if the regions, either gained or lost from the primary genome, are enriched for genes with particular functions. We used the genome annotation from Gounot et al. (2020) and checked for Gene-Ontology (GO) terms in regions that have gained or lost a copy of the primary genome. Additionally, we checked regions that are different in ploidy between the beer and the wine 1 subpopulation. Both approaches however, revealed no enrichment (data not shown). Further, we focused on the set of 56 candidate genes described in Colomer et al. (2020) (Supplementary Table S4). These genes are associated with particular functions and phenotypes, such as maltose assimilation, ethanol production or sulphite tolerance and play important (positive and negative) roles in different industrial applications. We could not find a pattern in which particular gene groups are linked to regions that are characterized by a conserved number of genomic copies for the primary genome (Supplementary Figure S9). Either these patterns of conservation have been acquired through adaptive processes for which we could not find any proof, or alternatively they have occurred though random processes. As already seen for the polyploid wine 2 subpopulation, LOH events are shared among strains and, with LOH events at similar positions (*e.g.* DEBR0S1), hotspots for LOH might be involved. However, we have no evidence for these at this point. Further, we would conclude that a recent ancestry could be also an explanation for the underlying pattern of LOH events, but due to the different origins of isolation in space and time, we believe it is highly unlikely to argue that the observed and conserved pattern within the subpopulations is due to a recently shared common ancestry.

Overall, the three subpopulations with polyploid genomes coming from interspecific hybridization events are highly dynamic, where LOH events have caused conserved patterns of low genetic diversity regions within each subpopulation. How these variations, especially on gene level are finally expressed at a phenotypic level, will have to be the goal of following studies investigating the phenotypic landscape of the different subpopulations.

### Expanded mitochondrial genomes and large inversions for the teq/EtOH subpopulation

With the polyploid subpopulations having undergone massive and independent modifications of their nuclear genome, we checked if this also accounts for their mitochondrial (Mt) genome. For this, we generated *de novo* assemblies from shortread sequencing data. We were able to prepare single circularized scaffolds for 48 of the 71 strains (Supplementary Table S5). Overall, *de novo* assemblies revealed a size between 75.3 kb and 89 kb for all subpopulations, except for teq/EtOH subpopulation, which Mt genome size was above 100 kb (Figure 5A).

**Figure 5.**
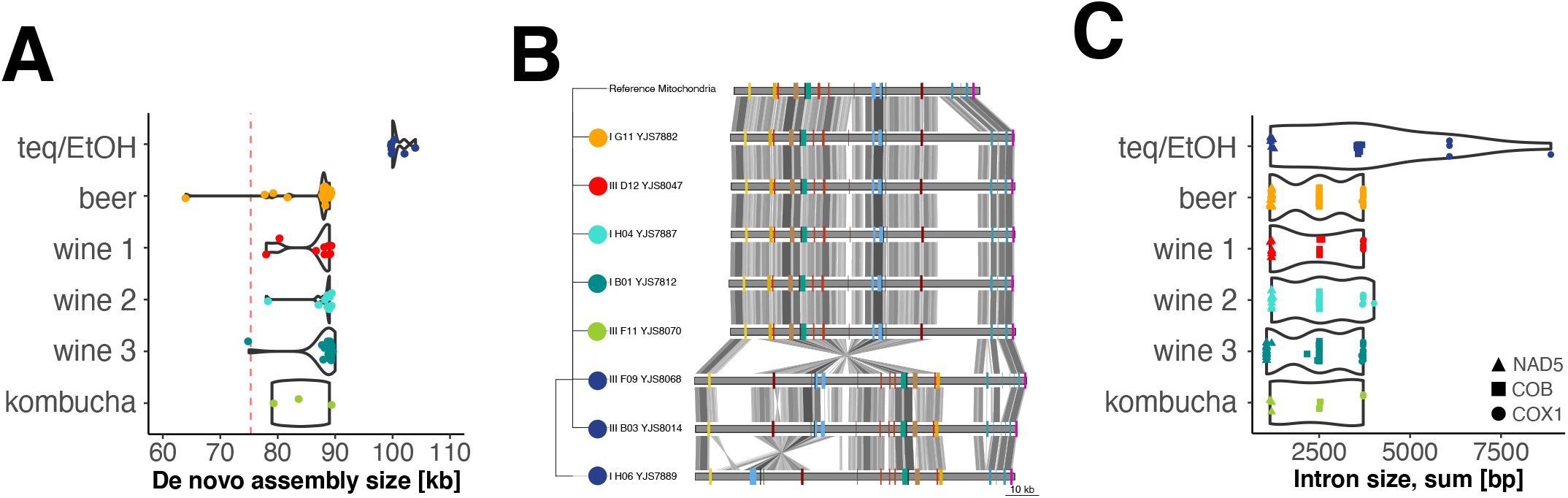
*De novo* assemblies for the mitochondrial genome reveal its expansion in all subpopulations and high level of reorganization for the teq/EtOH strains. **A** *De novo* assembly size difference. All subpopulations increased their mitochondrial genome compared to the reference (accession number GQ354526.1), from 75.3 kb to 88-90 kb. The teq/EtOH subpopulation stands out with *de novo* assembly sizes larger than 100 kb. Red dotted line: mitochondrial size of the reference genome. **B** Mitochondrial genome synteny. Synteny is conserved among subpopulations (compared to the reference), with the exception for the teq/EtOH subpopulation. Additionally, synteny within teq/EtOH was not congruent among strains. Here, the strain I_H06_YJS7889 harbours a large inversion. **C** Intron content contributes to genome expansion of teq/EtOH subpopulation. For teq/EtOH strains, the two protein coding genes *COB* and *COX1* have increased in intron content, while other subpopulations have the expected intron sizes compared to the reference genome (accession number GQ354526.1).

We then extracted gene positions and prepared a synteny approach to look for the organization of the different mitochondrial genomes. We can show that the teq/EtOH strains, apart from their extended sequence length, harbor several large inversions, while the organization of the mitochondria from the other subpopulation are aligned with the organization found for the reference (Figure 5B).

By calculating pairwise genetic distances from concatenated gene sequences, teq/EtOH strains show to be clearly distinct from all other strains (Supplementary Figure S10). Highest genetic distance of 3% was detected between two wine strains (I_B01_YJS7812 =wine 2; II_B03_YJS7914 =wine 3) with the teq/EtOH strains.

By looking at intronic content, we can show that two of the intron-carrying and proteincoding mitochondrial genes, *COB* and *COX1,* are partially involved in the overall size increase of the mitochondria in teq/EtOH strains. The other strains show no difference in intronic regions (Figure 5C).

## DISCUSSION

The *Brettanomyces bruxellensis* yeast species is known as harbouring subpopulations with various levels of ploidy (Avramova et al. 2018, Gounot et al. 2020, Colomer et al. 2020). For the first time, we provide a detailed insight into the complex genomic architecture of these polyploid subpopulations. Interestingly, we noticed that there is a high conservation of ploidy in each subpopulation and four of them, associated with three different ecological environments (tequila/ethanol production, wine making and beer brewing) are exclusively characterized by triploids.

Because polyploidy can be achieved in different ways (allopolyploidization or autopolyploidization), the final genomic composition might vary by distinct levels of intra-genomic information. At the same time, the intra-genomic variation will define the boundaries of genomic flexibility, and therefore drive evolution in almost unpredictable and different ways (Ng et al. 2012, Selmecki et al. 2015).

By using two different phasing strategies, we elucidated the genomic architecture of polyploid subpopulations of *B. bruxellensis* with various levels of intra-genomic variation. We highlighted that all six population harbour a primary genome irrespective of their ploidy, which is defined as the genetic variation that does not exceed the 1% compared to the reference genome *B. bruxellensis* (Fournier et al. 2017). This is lower but in accordance with previous papers elucidating the genetic variation of *B. bruxellensis,* since they did not phase the genomes into distinct haplotypes (1.2%; Gounot et al. 2020). Further, we can show the existence of three allopolyploid subpopulations (teq/EtOH, beer, and wine 1) with an acquired genomic copy with a genetic divergence of about 3% compared to the reference genome. They clearly exceed the average intra-genomic variation of the primary genome, undermining the occurrence of interspecific hybridization events in these subpopulations. The known sister species within the same genus are rejected as donors for the interspecific hybrids due to too high genetic divergence of at least 23% (Roach and Borneman 2020).

We further highlight the fact that to our knowledge, the *B. bruxellensis* species is one (or the) rare case, in which these different scenarios, respectively allo- and autopolyploidy, can be observed in closely related subpopulations. We observed different trajectories for strains not only associated with different environments (teq/EtOH, beer and wine) but also associated with the same environment, while being part of a genetically distinct cluster. The “wine”-associated strains fall in three genetically diverged subpopulation. With the two subpopulations wine 1 and wine 2 being triploid compared to wine 3 (diploid), only wine 1 has acquired a third genomic copy from interspecific hybridization, while wine 2 has solely genetically similar haplotypes.

Different trajectories of polyploidizations in nature were mostly studied (and observed) in plants, which give no clues about the importance as well as prevalence of polyploidization and their trajectories in animal or fungal systems (Leggatt and Iwama 2003; Gregory and Mable 2005; Barker et al. 2015). At the same time, these mechanisms, when observed and studied in extant polyploids, have mostly, when not exclusively been determined between species systems, rather than within. We highlight that future studies, especially in the animal and fungi kingdom are required, which screen individuals on a large scale, to study prevalence and trajectories of polyploids across ecologically diverged, naturally occurring subpopulations. Indications that polyploidy could be a more common state were shown by two recent studies, which genotyped more than 1,000 individuals from the *S. cerevisiae* and *B. bruxellensis* yeast species, with a prevalence of polyploids of 11,4% and 54%, respectively (Avramova et al. 2018; Peter at al. 2018).

Finally, we speculate that the different trajectories of polyploids in the subpopulations of *B. bruxellensis* are linked to the adaptation to the different anthropized environments. Polyploids in general have been given a lot of attention in the context of adaptivity and diversification, in which many extant species originate from ancient polyploid states (Seoighe and Wolfe 1998; Gordon et al. 2009; Gertein and Otto 2009; Marcet-Houben and Gabaldon 2015). While a polyploid state itself can allow adaptability, it is often seen as a transient state, which is followed by massive modifications thereafter to cope with genetic incompatibilities and to regain fertility in the long term. Evidence for this process has been gained through the detection of paralogous gene sets with different historical trajectories in many naturally diploid taxa, undermining the process of genomic modifications after polyploidization. With a prevalence of 54% polyploids, plus evidence for three independent interspecific hybridization events, polyploidy is very abundant and most likely does not underlie random effects for *B. bruxellensis.* The genomes of the allo- and autopolyploid subpopulations are characterized by massive genomic modifications, which have established conserved pattern of rearranged blocks. These underline on the one hand the independent acquisition of genetically diverse genomic copies for the allopolyploid subpopulations, but most likely reflect the regain of fitness and overcome of genomic incompatibilities, while being able to adapt to harsh and changing conditions in their anthropized environments. The high tolerance against sulphur dioxide for example, a treatment to prevent wine fermentation from spoilage by *B. bruxellensis*, could be the response of this yeast to the recently increased usage of this agent from the industry, and have been mostly observed for the wine 1 subpopulation (Avramova et al. 2019). Our study clearly highlights for the first time the coexistence of a large repertoire of evolution punctuated by various independent polyploidization events within a species and addresses the need to further resolve the genomic architecture of polyploid species complexes from diverse ecological settings.

## Material and Methods

### Selection of strains and DNA extraction

For this study, we focused on a subset of 71 strains of *Brettanomyces bruxellensis.* These strains are part of the collection of 1,500 strains (Avramova et al. 2018), which was previously analyzed using microsatellites and/or partially with whole genome sequencing data (Gounot et al. 2020). The 71 strains were selected to represent the different clades of *B. bruxellensis* in terms of genetic diversity, ecological origin (origin of isolation) and variation in ploidy (Supplementary Table S1). Additional to these 71 *B. bruxellensis* strains, a representative of four sister species (*B. anomala, B. custersianus, B. nanus, B. acidodurans*) as well as the type strain of *B. bruxellensis* were selected for this study (Supplementary Table S3).

The DNA of the 76 strains was extracted from 20ml cultures (single colony, 48h growth at 25°C) using the QIAGEN Genomic-tip 100/G kit (Hilden, Germany) with the recommended manufacture’s genomic DNA buffer set. The manufacture’s protocol was followed as recommended and final DNA was eluded in 100-200*μ*l water. DNA was quantified with the broad-range or high-sensitivity DNA quantification kit from Qubit (Thermo Fischer Scientific, Waltham, USA) with the use of the automated plate reading platform from TECAN (Männedorf, Switzerland). Genomic DNA was migrated on a 1.5% agarose gel to check for degradation.

### Library preparation and sequencing

The kit NEBNext® Ultra™ II DNA Library Prep Kit (Ipswich, USA) for Illumina® (San Diego, USA) was used for library preparation. The dual-barcoding strategy was applied and samples were sequenced on two lanes of NextSeq (Illumina®) at the European Molecular Laboratory (EMBL) in Heidelberg, Germany. The strategy of sequencing was 75 paired-end (75PE) and sequences from two independent sequencing lanes were concatenated prior to any analysis.

For the long-read sequencing we used the Oxford Nanopore Technology (Oxford, UK). Libraries for sequencing using the MinION and were prepared as described in (Istace et al. 2017) using the Ligation Sequencing Kit SQK-LSK109. We barcoded strains with the Native Barcoding Expansion 1-12 (EXP-NBD104) to multiplex up to 12 samples per sequencing reaction.

## Data analyses: long reads (MinION)

### Base-calling, de-multiplexing and adapter trimming

Raw sequencing reads were processed as described in (Fournier et al. 2017). Briefly, base-calling and de-multiplexing was done with guppy (https://nanoporetech.com/). Adapters were trimmed with Porechop (Porechop GitHub Repository https://github.com/rrwick/Porechop).

### Separating reads with different degrees of genetic variation to the reference genome

We distinguished reads depending on their genetic divergence to the reference genome of *B. bruxellensis* (Fournier et al. 2017). For this, long reads of each sample were first aligned to this sequence using MiniMap (Li 2018). We separated reads into two groups based on their number of SNPs/kb. Here, reads comprising less than ten variants per kb were assigned to the low intra-genomic variation cluster and reads with more than 14 variants per kb to the high intra-genomic variation cluster. Reads containing between ten to 14 variants per kb were ignored to avoid any errors of wrong-assignment, which could strongly impact *de novo* genome assemblies.

### Calculation of coverage for the low and high intra-genomic variation cluster

The contribution to the overall coverage (total coverage) was calculated for the reads that clustered in the low intra-genomic variation cluster and the high intra-genomic variation cluster. This was done in 10kb non-sliding windows and used as a measurement of the average ploidy per strain. As an example, if the overall coverage for a certain region was calculated to be 60x (from reads with low and high intra-genomic variation), then a coverage of 40x for the reads with low intra-genomic variation and 20x for the reads with high intra-genomic variation would assume a triploid state at this locus, with a ratio of genomic copies of 2:1. This method was adapted to estimate different potential levels of ploidy (2n to 5n).

### Phasing the polyploid genomes of the wine 2 subpopulation

We phased six polyploid genomes of the wine 2 subpopulation with the nPhase pipeline as described in (Abou Saada et al. 2020). For this, short and long reads were aligned to the *B. bruxellensis* reference sequence and phased by the nPhase algorithm using default parameters.

To generate pairwise divergence plots, we cross-referenced two of the files output by nPhase, located in the Phased folder: (1) the *.clusterReadNames.tsv file, which contains the list of reads that comprise each cluster and (2) the *.variants.tsv file, which contains the list of heterozygous SNPs associated with each predicted haplotig. By combining the information in both files we were able to calculate the similarity between predicted haplotypes in 10kb windows.

In regions that have only two predicted haplotypes we have only one value, but in regions that have more than three predicted haplotypes we only kept the three longest clusters and generated three similarity values through pairwise comparison (used for potting maximal genetic distances between haplotypes).

### *De novo* assemblies

Prior to the *de novo* assemblies, files containing the raw reads (respectively with low or high intragenomic variation to the reference genome) were corrected and cleaned using Canu -correct v.1.7 (Koren et al. 2017). *De novo* assemblies were performed with SMARTdenovo (Liu et al. 2020) and the parameters -J 1000 -c 1.

### Collinearity and pairwise genetic identity of de novo assemblies

Collinearity between *de novo* assemblies of *B. brettanomyces* strains was checked using Mummer v.3 and the following parameters nucmer --mum -l 200. To check for collinearity between different species, we lowered the values and stringency to --mum -l 20 -c 30 -b 100.

## Data analyses: short reads

### Genome-wide phylogeny and estimation of ploidy

Raw sequencing reads (not separated short reads) were aligned to the reference genome *B. bruxellensis* (Fournier 2017) using BWA v0.7.17 (Li and Durbin 2009) with the default settings (mem algorithm). File format conversions, the sorting and indexing of samples were done with Samtools v.1.9 (Li et al 2009). Variant calling was done using the Genome Analysis Tool GATK v4.1 (McKenna et al. 2010). The data from the variant calling in GATK was filtered and processed with VCFtools (Danecek et al. 2011) and BCFtools v1.9 (Li et al. 2009). Respectively, we filtered out any indels, kept only variants with a minimum coverage of 11 reads/site, removed individuals with more than 50% of missing data and reduced the data. The information of the Allele Balance for the Heterozygous sites (ABHet) was used to calculate the average allele frequencies in 10kb windows (non-sliding) in R v.3.3.3 (R Core Team 2019). Phylogenetic Neighbor-Joining trees were performed with the R packages seqinr (Charif and Lobry 2007) and phangorn (Schliep 2011) using the substitution model JC69. The final trees were plotted with Figtree v.1.4.3 (Rambaut 2009).

### Genomic-copy specific alignments

A competitive mapping approach was used to distinguish short reads that represent the low or high intra-genomic variation. For this, the short reads of the 40 strains from the three polyploid subpopulations with low and high intra-genomic variation (teq/EtOH, beer, wine 1) were aligned to clade-specific reference genomes. These reference genomes were concatenated *de novo* assemblies, respectively prepared from low and high intra-genomic variation (long-read data) to the reference genome *B. bruxellensis*. These clade-specific reference genomes came from the polyploid strains YJS7895 (beer), YJS8039 (wine 1) and YJS7890 (teq/EtOH). Finally, to have all reads aligned to the same reference genome and to perform comparative genomic analyses, the reads from the competitive mapping approach, which either mapped on the scaffolds from the low or the high intra-genomic *de novo* assemblies, were mapped back to the reference genome of *B. bruxellensis* (Fournier et al. 2017). The 31 strains, which did not show any signals of polyploidy (wine 3, kombucha) or high intra-genomic variation (wine 2) were mapped directly to the *B. bruxellensis* reference genome. In this way, all strains were ultimately aligned to the same reference facilitating the direct comparison of genetic variation. Alignments, file conversions, file sorting, file indexing and the calculation of coverage in 10 kb windows were done as described above.

### Principal Component Analysis and phylogenetic analysis

Variant calling and filtering were done as described above. The program Adegenet v2.1.0 (Jombart 2008) was used to perform the Principal Component Analysis (PCA). Phylogenetic trees were generated and plotted as described above.

### Pairwise distances

Samtools v1.9 and Bcftools v1.9 (Li et al. 2009) were used to calculate the genotype likelihood from the bam-formatted alignment files, to call variants and to create single fasta files for each individual strain. Genetic distances were calculated in 50 kb windows in R with the package phangorn (Schliep 2011; substitution model “JC69”) and then averaged per individual.

### Detection of regions underlying the variation in copy numbers

Variation in copies of the low intra-genetic variation along the polyploid genomes of the 40 allopolyploid strains was calculated in 10 kb windows from the ratio of the coverage of the primary genome to the total coverage. Ploidy levels were categorized as described above. Plots were generated with the R package ggbio (Yin et al. 2012).

### Gene ontology enrichment and candidate gene approach

Gene ontology enrichment were done using a reference list of 2,274 annotated genes with orthologous genes in *S. cerevisiae* (Gounot et al. 2020). Genes were considered to be enriched (and included in the analyses) when at least 50% of the strains in the two groups beer and wine 1 contained the same copy number of the gene from the primary genome. The group Tequila/Ethanol was not included in this analysis due to the low number of strains and the structure of two sub-groups. GO enrichment analysis was performed using with the program Gorilla (Eden et al. 2009). Candidate genes were chosen from Colomer et al. (2020) and plotted according to their position in the genome. Here, we used the gene positions from the genome annotation of Gounot et al. (2020).

### Analysis of mitochondrial genomes

The mitochondrial *de novo* genome assemblies were constructed with a pipeline derived from Tao et al. (2019). Illumina reads were down sampled to sets of 600,000, 800,000 and 1,000,000 paired-end reads with seqtk (https://github.com/lh3/seqtk) and *de novo* assemblies were constructed for each dataset with A5-miseq (Coil et al. 2015). Mitochondrial contigs were identified through similarity searches to the *B. bruxellensis* mitochondrial reference sequence (accession number GQ354526.1) and for each strain, a representative assembly was selected based on the number of contigs and their length. The one-contig assemblies were subjected to circlator (Hunt et al. 2015) for circularization and a custom python script was used to set the starting position of the sequence to that of the reference genome.

Synteny between strains from each group were compared in Mauve (Darling et al. 2004) using the MauveAligner algorithm with the default parameters. Synteny was graphically displayed using the R package genoPlotR (Guy et al. 2010) and the coordinates of introns and exons. Phylogenetic trees of mitochondrial genes were prepared as described above. The concatenated tree based on eight protein coding genes, for which we had genetic information of 70 strains (*ATP6, ATP8, COX2, COX3, NAD1, NAD3, NAD4, NAD4L, NAD6).* Genetic distances were computed from the concatenated genes alignments with the substitution model JC69.

## Availability of data

Illumina and Oxford Nanopore data for the 71 *Brettanomyces bruxellensis* isolates are available under the study accession number PRJEB41126.

Oxford Nanopore sequencing data for the *B. anomala, B. nanus, B. custerianus* and *B. acidodurans* species is available under the study accession number PRJEB41125.

## Acknowledgements

This work was supported by the Agence Nationale de la Recherche (ANR-18-CE20-0003-02 and ANR-18-CE12-0013-02) and the European Research Council (ERC Consolidator Grant 772505). J.S. is a Fellow of the University of Strasbourg Institute for Advanced Study (USIAS) and a member of the Institut Universitaire de France.

